## Supplemental Materials for "Simultaneous enhancement of multiple functional properties using evolution-informed protein design"

(1) Department of Systems Biology, Harvard Medical School, Boston, MA, USA
(2) Program in Biomedical Informatics, Harvard Medical School, Boston, MA, USA
(3) Institute for Protein Innovation, Boston, Massachusetts, Boston, MA, USA
(4) Division of Hematology/Oncology, Boston Children's Hospital, Harvard Medical School; Boston, MA, USA
(5) current address: AI Proteins; Boston, MA, USA
(6) Department of Data Sciences, Dana-Farber Cancer Institute, Boston, MA, USA
(7) current address: Research Institute of Molecular Pathology (IMP), Vienna BioCenter (VBC), Campus-Vienna-Biocenter 1, 1030 Vienna, Austria
(8) School of Biochemistry and Immunology, Trinity College Dublin, Dublin 2, Ireland
(9) Division of Newborn Medicine, Boston Children’s Hospital, Boston, MA, USA
(10) Selux Diagnostics, Inc., 56 Roland Street, Charlestown, MA, USA

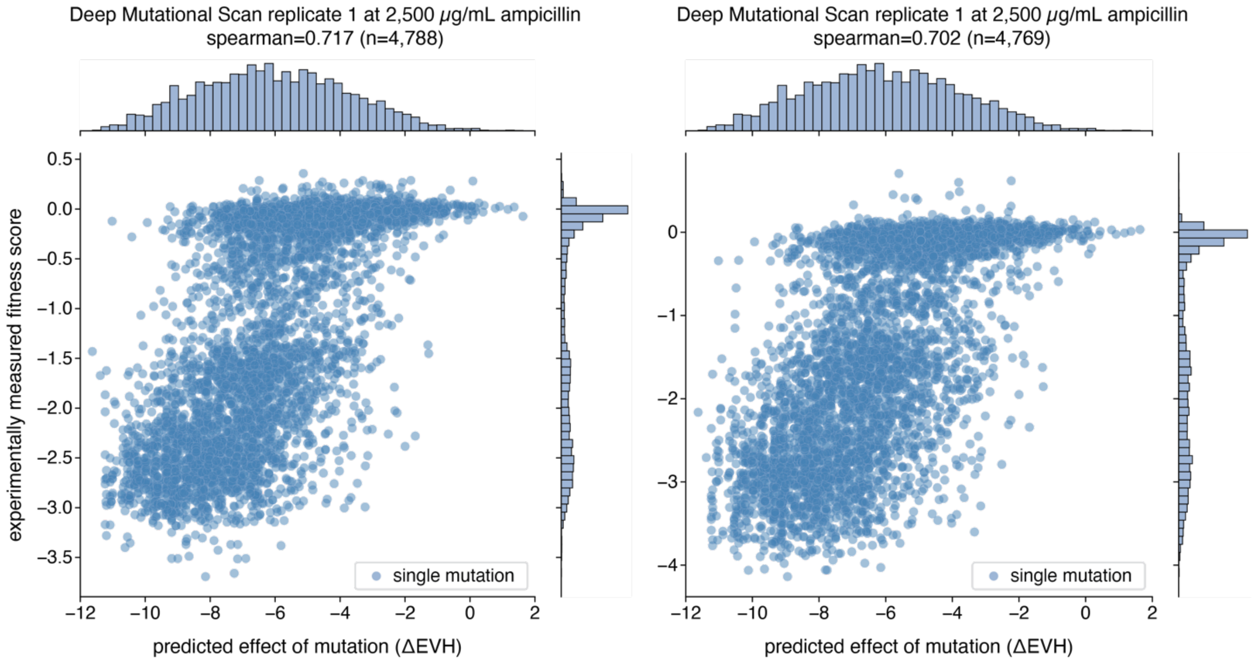


**Supplemental Figure 1: Relationship between predicted fitness score and experimentally-determined fitness effect of individual mutations from a published deep mutational scan [1]**. Each blue dot represents a single point mutation of which there are 4,788 possible mutations over 252 positions aligned in the multiple sequence alignment used for model generation and fitness prediction. The predicted fitness effect (ΔEVH) on the x-axis is the predicted fitness of the point mutant minus the predicted fitness of WT TEM-1. The experimentally measured fitness score on the y-axis is defined in Stiffler et al. [1]. **Left**: Experimental replicate 1, which quantified all possible mutations (n=4,788 with spearman=0.717). **Right**: Experimental replicate 2, which quantified all mutations in 251 positions (n=4,769 with spearman=0.702).

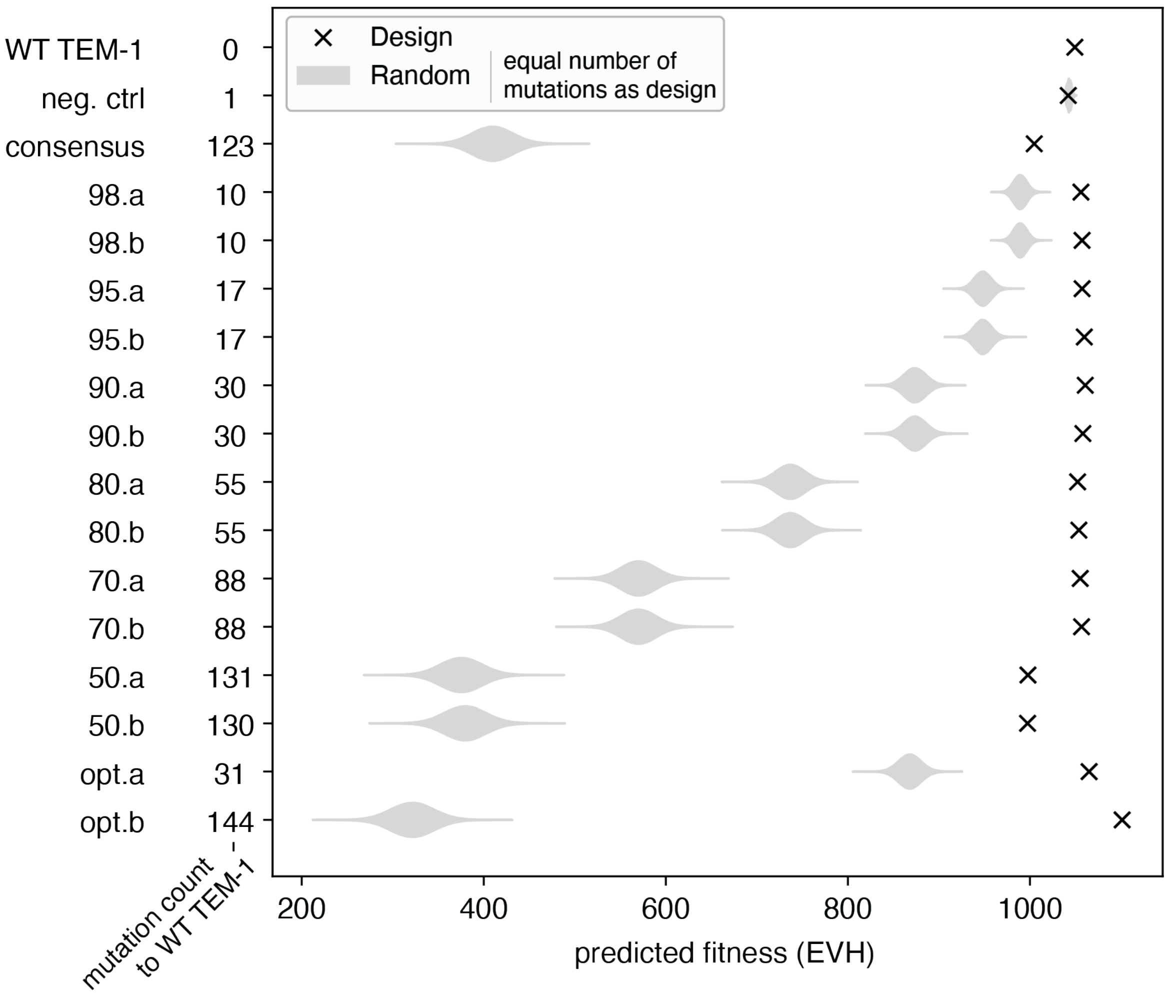


**Supplemental Figure 2: Comparison of each designs' predicted fitness with randomly generated sequences.** For each design, 1 million random protein sequences were generated on the WT TEM-1 sequence background with an equal number of mutations as the design. The predicted fitness of each design is indicated as an "X" and the distribution of the random sequences is shown as a violin plot. In general, the designs had a much higher predicted fitness than random variants.

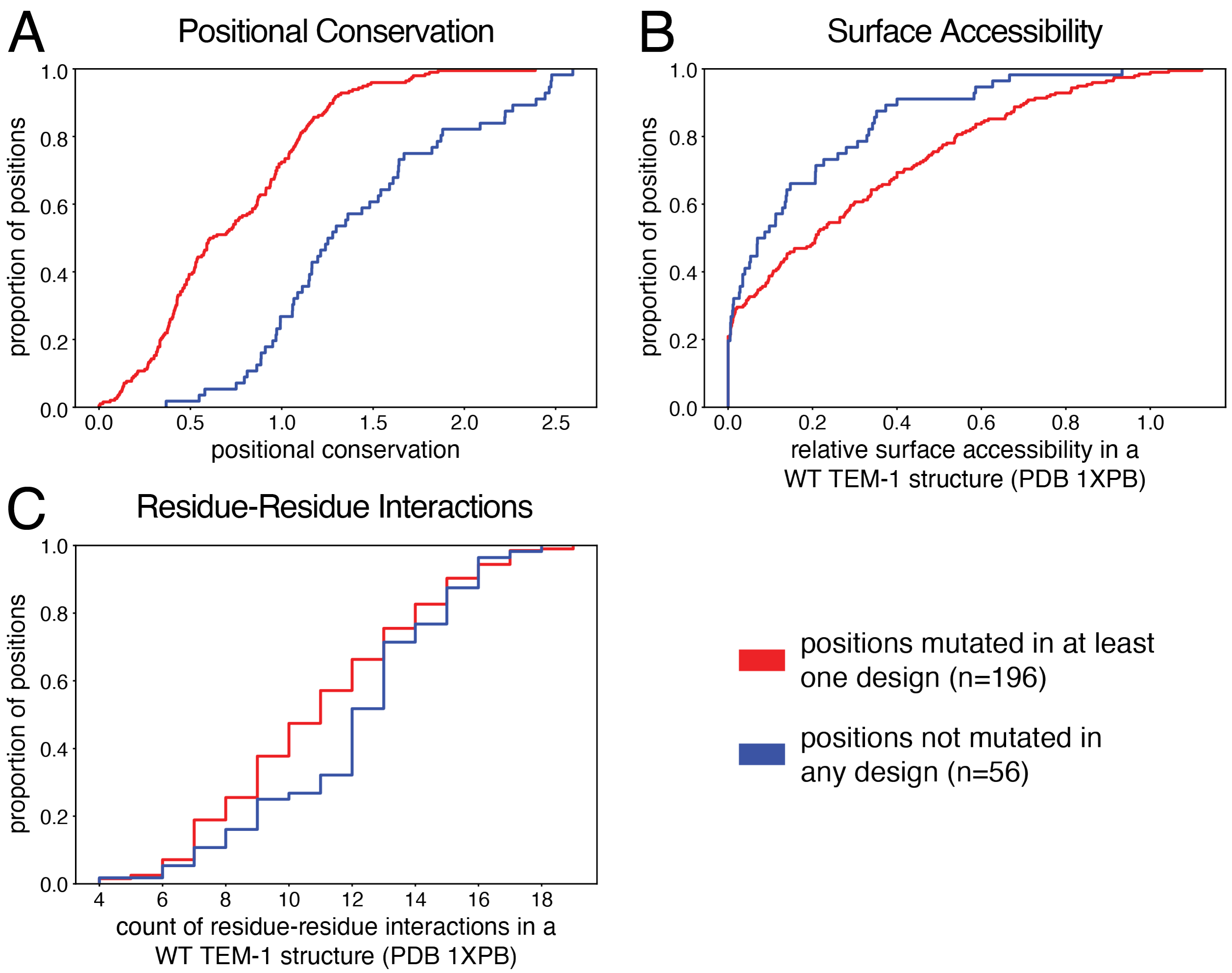


**Supplemental Figure 3: Properties of positions mutated in all of the designs compared to non-mutated positions.** Each panel contains two cumulative distribution functions (CDFs): positions mutated in one or more designs (red) and positions not mutated in any design (blue). The y-axis is the proportion of positions that are equal or less than the value of the property shown on the x-axis. **(A)** CDFs of positional conservation (maximum shannon entropy of all positions - shannon entropy at the positions). Positions mutated in the designs were less likely to be conserved than positions not mutated. **(B)** CDFs of relative surface accessibility from DSSP analysis (relative_acc output) of a published WT TEM-1 structure (PDB 1XPB). Residues mutated in the designs were less likely to be surface accessible than non-mutated residues. **(C)** CDFs of residue-residue interaction counts from a published WT TEM-1 structure (PDB 1XPB). An interaction is defined as having any atom in one residue be within 5 angstroms of any atom in the other residue. Residues mutated in the designs generally had fewer interactions than non-mutated residues.

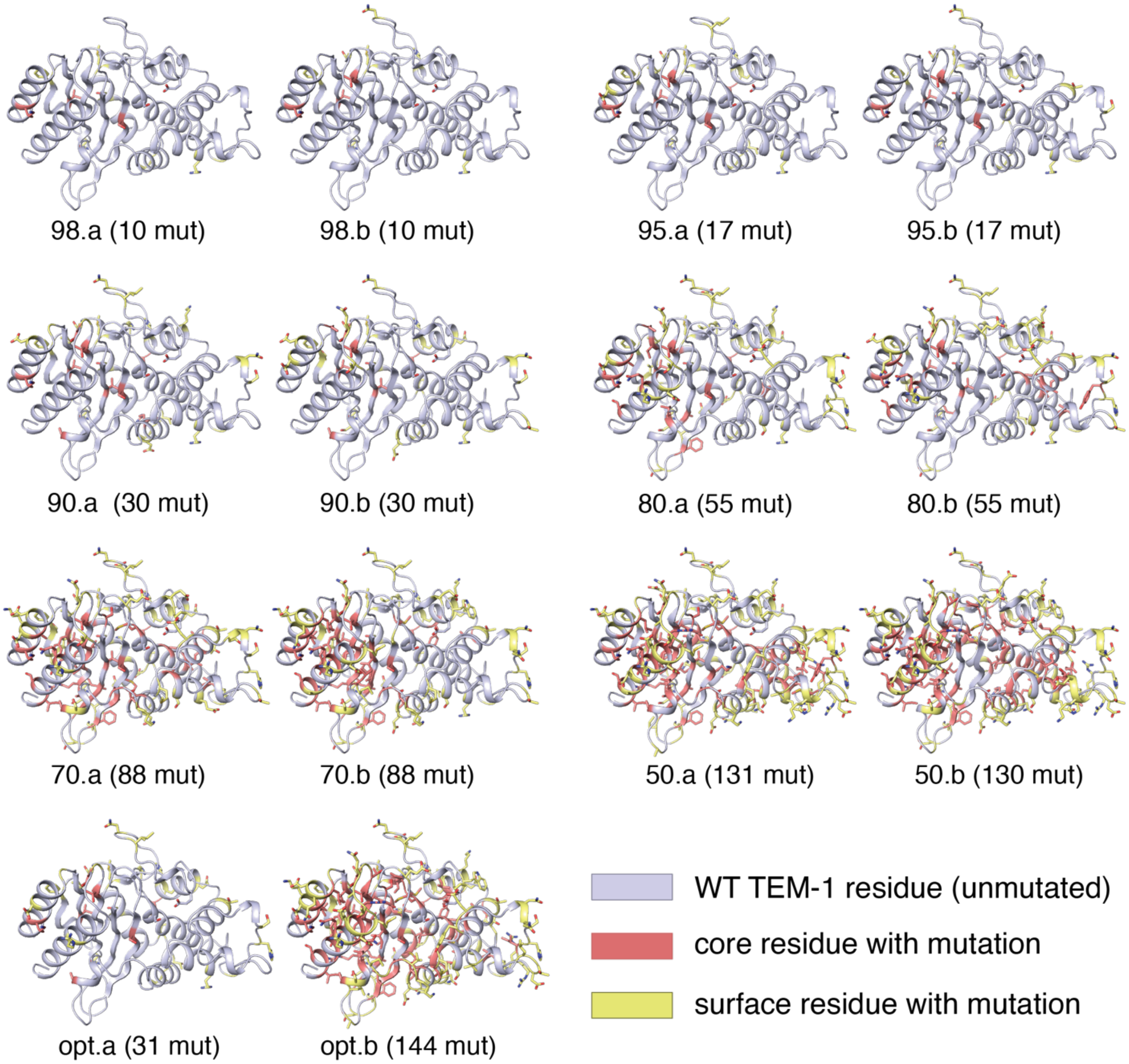


**Supplemental Figure 4: Structural representation showing the positions mutated in each design on a published WT TEM-1 structure (PDB 1XPB).** The position of each design mutation is shown with stick representation of the wild type amino acid, and is colored according to relative solvent accessibility. Yellow = surface residues (relative solvent accessibility of >= 20%). Red = core residue (relative solvent accessibility of < 20%). Unmutated positions are in cartoon representation in silver.

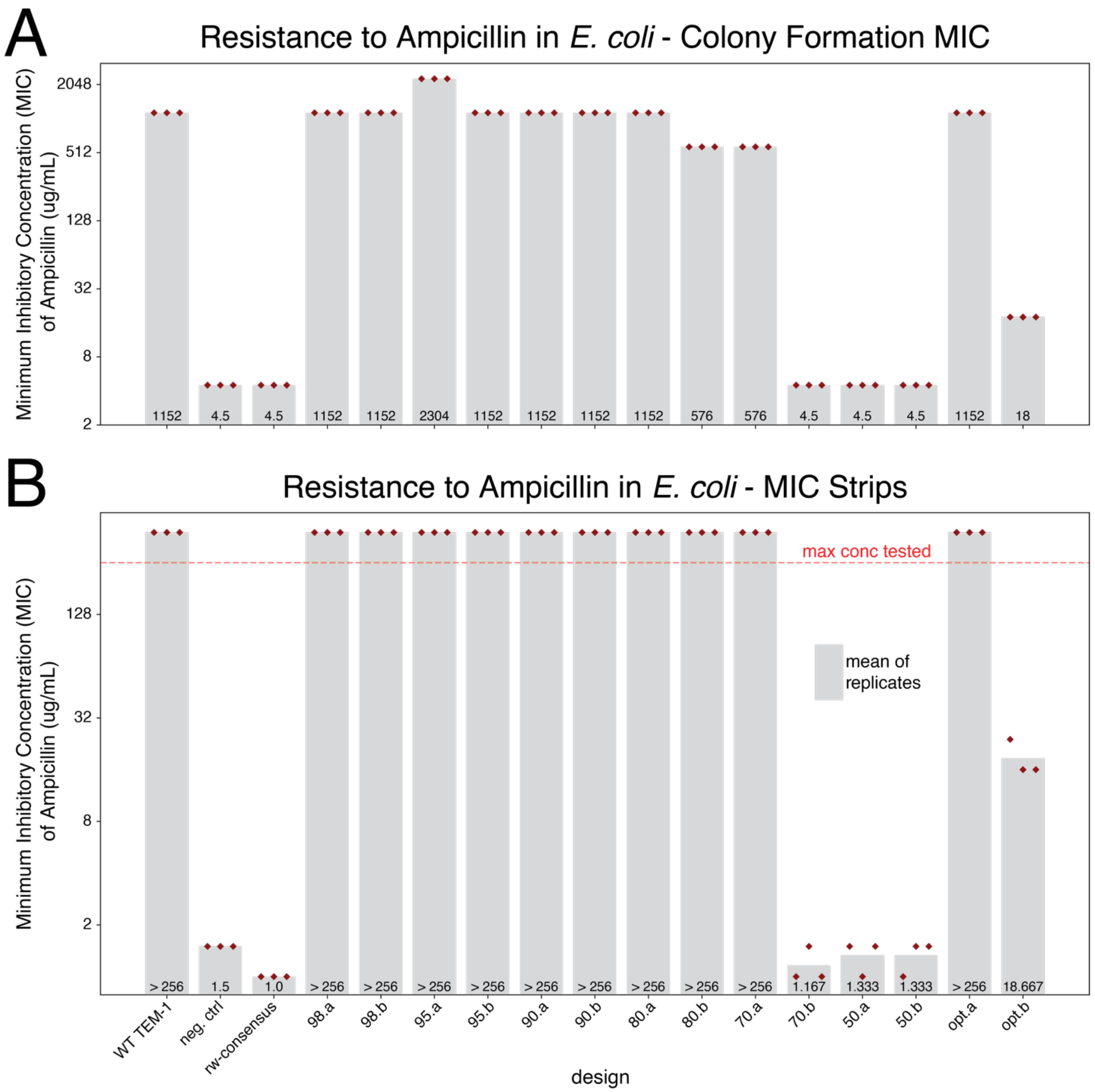


**Supplemental Figure 5: Ability of designs to confer resistance to ampicillin in *E. coli*.** In addition to the broth microdilution assay in the main text (Figure 3), two independent assays were used to measure minimum inhibitory concentration (MIC) of the canonical β-lactam substrate ampicillin. Resistance to ampicillin as assessed by **(A)** ability to form colonies across a serial dilution of ampicillin in MH agar, and **(B)** a MIC strip assay (Liofilchem). The maximum concentration of ampicillin tested, and therefore maximum MIC resolution, in the MIC strip assay in (B) was 256 μg/mL (indicated in red). The relative MIC between sequences was largely in agreement for all three resistance assays (broth microdilution, colony formation MIC, and MIC strips).

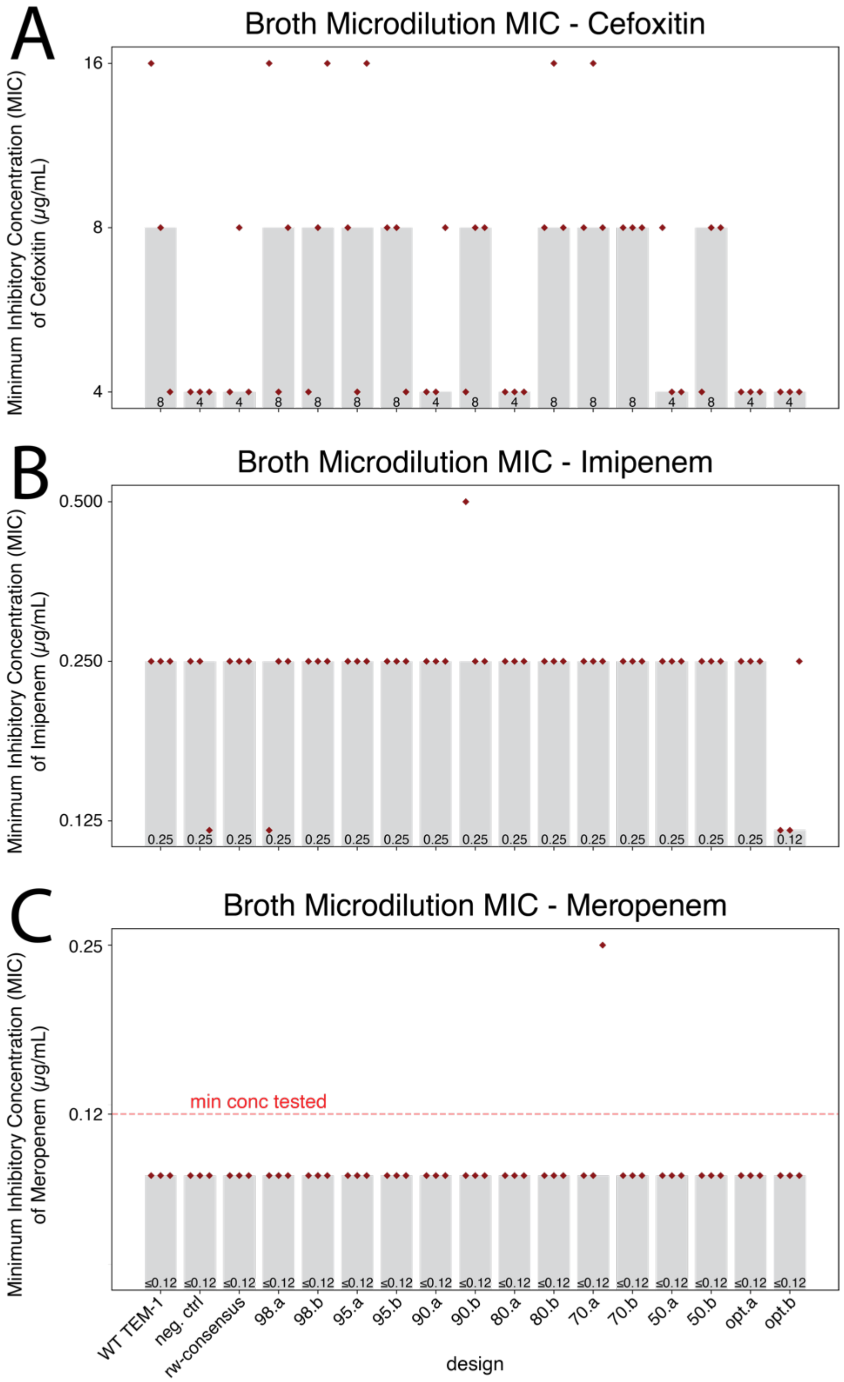


**Supplemental Figure 6: For some β-lactam antibiotics, the designs showed no consistent differences in their ability to confer resistance with WT TEM-1 or the negative controls.** Minimum inhibitory concentration (MIC) in *E. coli* was determined by a Clinical and Laboratory Standards Institute (CLSI) broth microdilution assay. The aggregated MIC calls (gray bars, see Methods) summarize three individual replicate experiments. (A) Resistance to Cefoxitin, a second-generation cephamycin β-lactam antibiotic. (B) Resistance to Imipenem and (C) meropenem, both of which are members of the carbapenem class of β-lactam antibiotics.

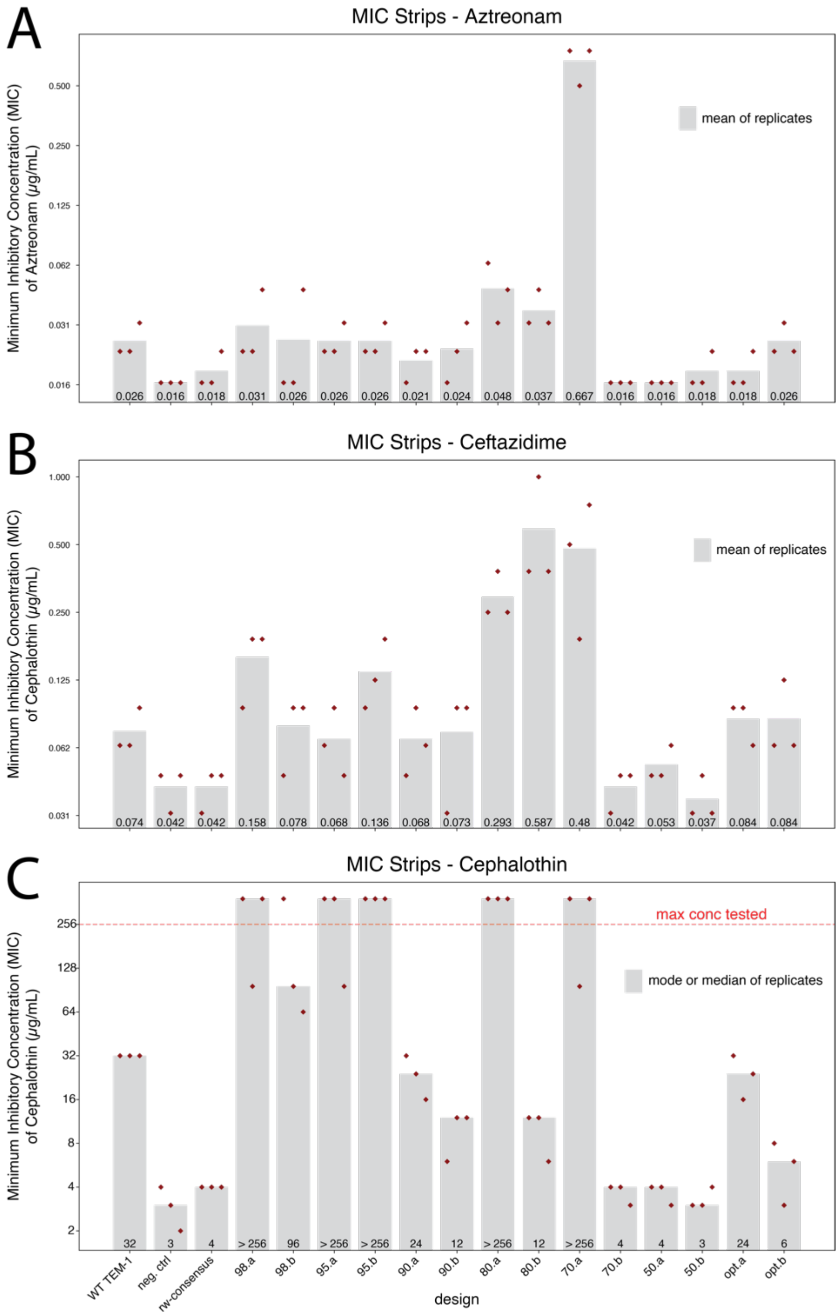


**Supplemental Figure 7: Ability of designs to confer resistance to β-lactam antibiotics aztreonam, ceftazidime, and cephalothin in *E. coli*.** Minimum inhibitory concentration (MIC) in *E. coli* was determined by a MIC strip assay (Liofilchem). (A) MIC of aztreonam. Gray bar: mean of three replicates. (B) MIC of ceftazidime. Gray bar: mean of three replicates. (C) MIC of cephalothin. For cephalothin, several of the replicates exceeded the maximum concentration tested (256 µg/mL), and therefore the gray bars are either the mode of the replicates or, if there was no mode, the median of the three replicates. The relative MIC between sequences was largely in agreement with the broth microdilution resistance assays in the main text (Figure 4).

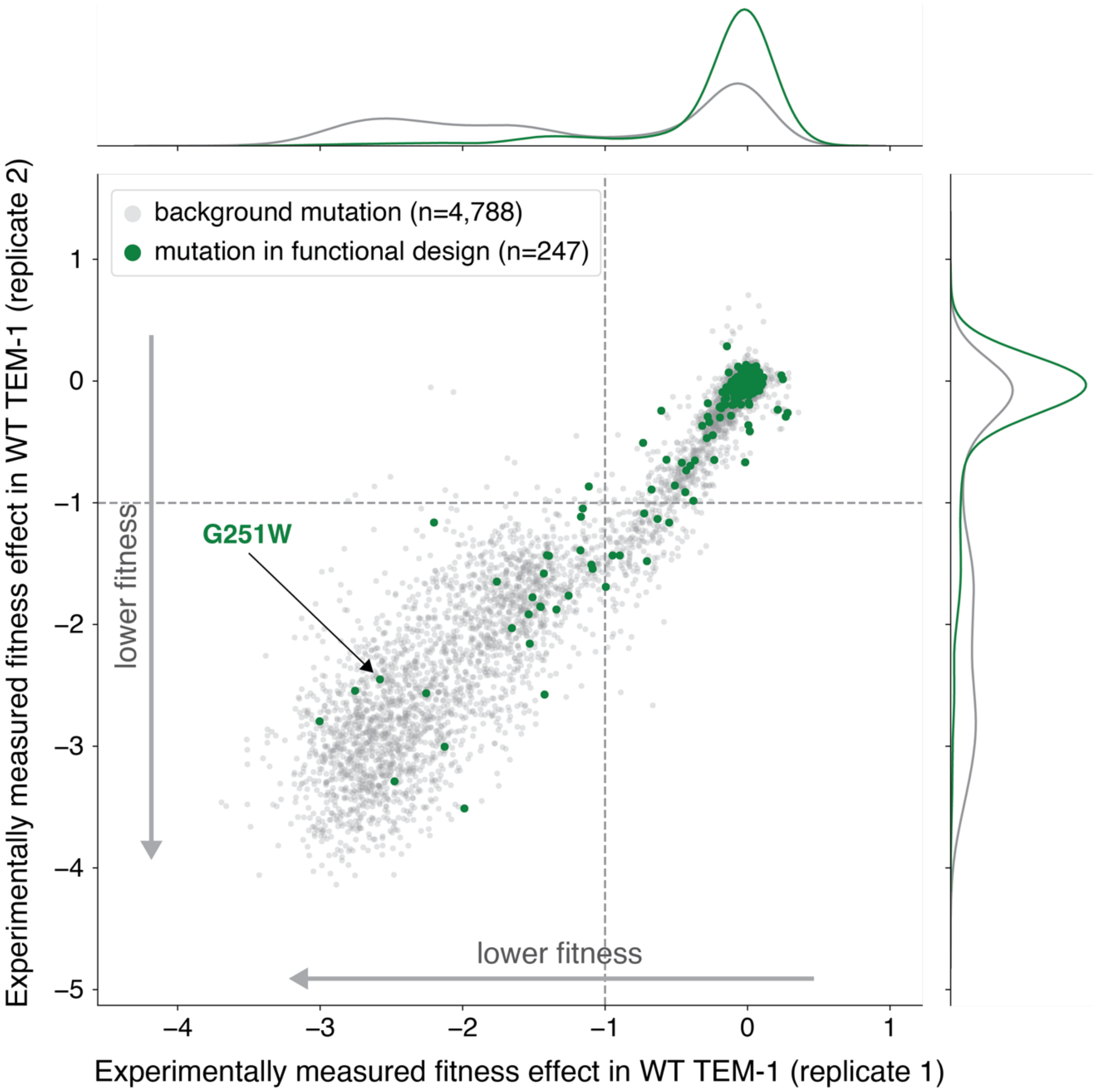


**Supplementary Figure 8: Functional design variants contain mutations that negatively affect fitness in WT TEM-1 in a previously published deep mutational scan by Stiffler et al. [1].** Comparison of aggregated amino acid changes from every functional design to the experimentally-determined fitness effect all possible point mutations in the WT TEM-1 sequence background (modeled polypeptide length of 252 multiplied by 19 possible substitutions). Each dot represents a single amino acid change, and axes quantitate the fitness effect at 2,500 µg/mL ampicillin (x-axis = replicate 1, y-axis = replicate 2). Dotted lines indicate a score of less than -1, which conceptually equates to a 10-fold decrease in fitness at a given ampicillin concentration relative to WT TEM-1. Several design variants (80.a, 80.b, 70.a, and opt.a) contain mutations that negatively affect fitness in isolation in WT TEM-1, i.e., have mutations that are represented in the lower left quadrant. Marginal distributions of the data are shown using kernel density estimation (KDE) on the top (replicate 1) and the right (replicate 2) of the scatter plots. [Colors] Gray: all point mutations for positions aligned in the β-lactamase multiple sequence alignment. Green: amino acid changes that occur in at least one functional design.

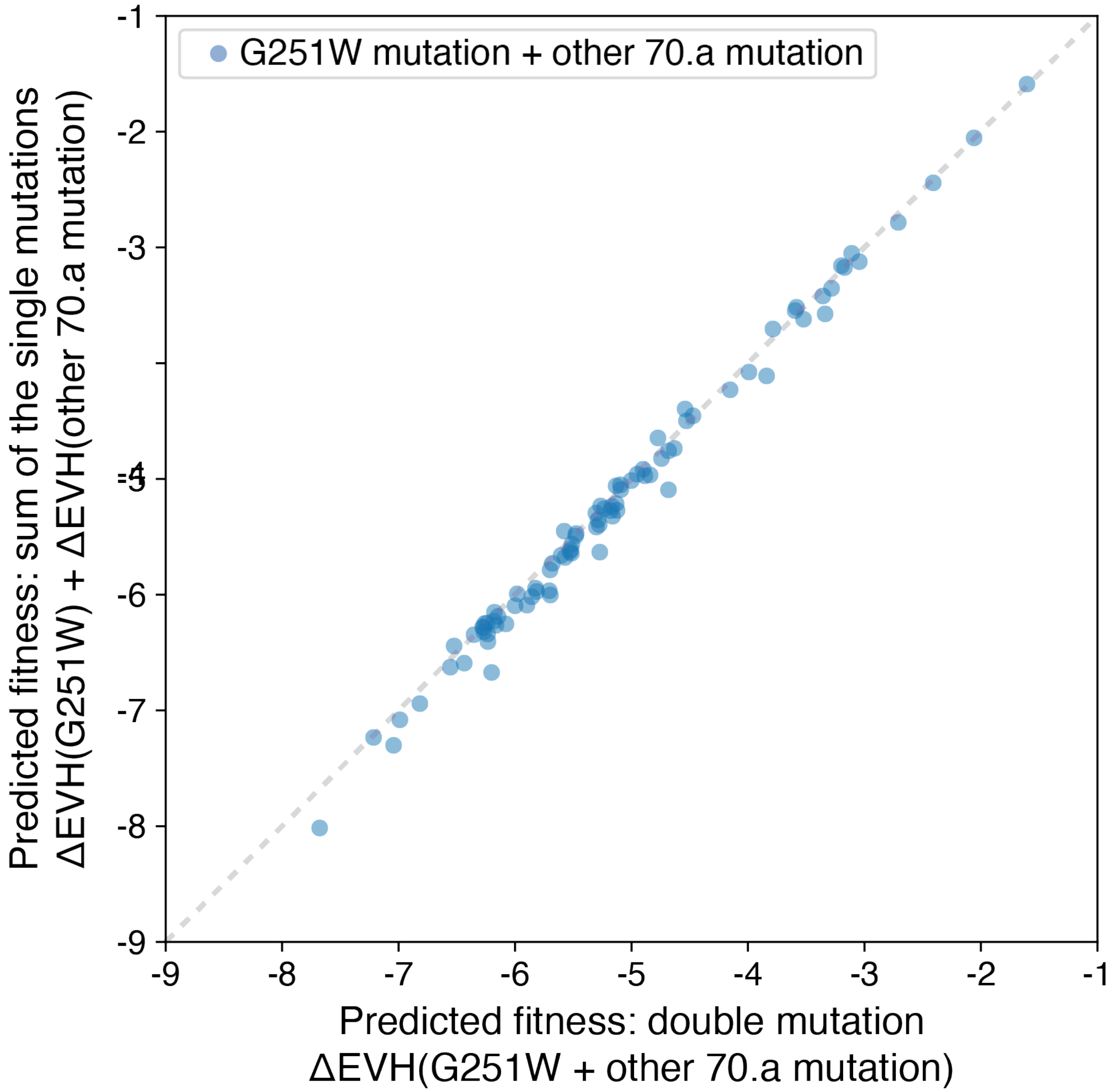


**Supplemental Figure 9: Comparison of the predicted fitness of double mutations versus the sum of the predicted fitness of single mutations for G251W with other 70.a mutations.** Each blue dot is a single 70.a mutation combined with the G251W mutation as a double mutation (x-axis) or the sum of the single mutations (y-axis). The similarity between these fitness predictions (Pearson $r$ = 0.996) indicates a lack of epistasis between G251W and the other 70.a mutations.

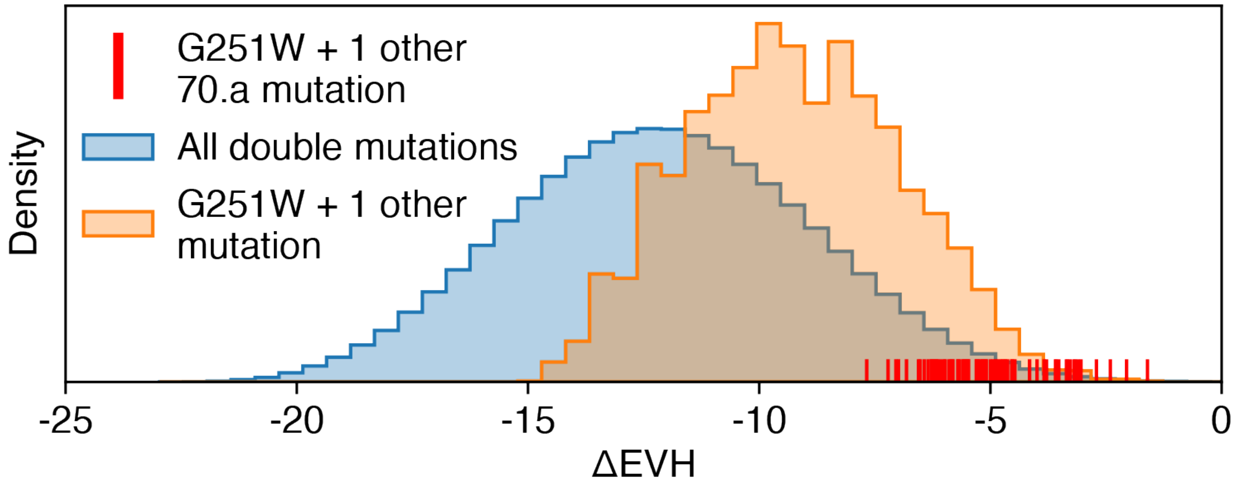


**Supplemental Figure 10: Comparison of the predicted fitness changes of double mutations in the WT TEM-1 sequence background with and without the G251W mutation.** The change in predicted fitness of pairs of mutations is calculated as the EVH of the double mutants minus the EVH of WT TEM-1. Blue distribution: all possible double mutations in WT TEM-1. Orange distribution: all double mutations that contain G251W. Red residual dashes show all double mutations with G251W and each 70.a mutation (n=87 doubles). The double mutations containing G251W are generally higher than random double mutations (orange versus blue), and individual mutations in 70.a when combined with G251W are on the right side of both distributions indicating they are amongst the most beneficial of double mutations.

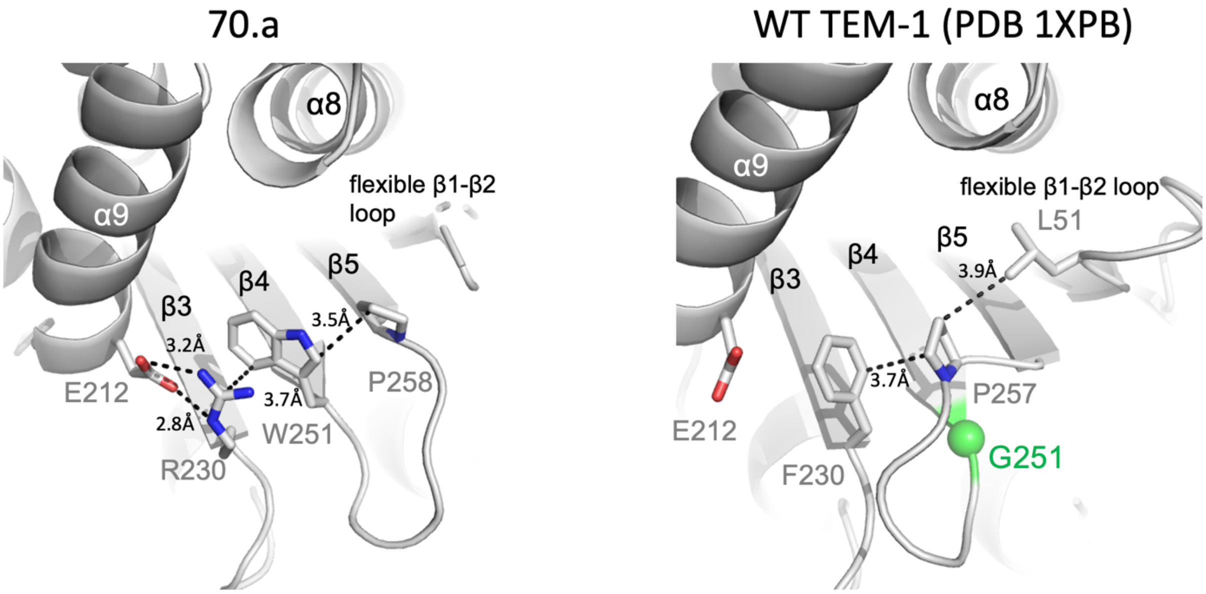


**Supplemental Figure 11: Alternative capping of the hydrophobic core of TEM-1 β-lactamase.** Focused visualization of position 251 in the crystal structure of the 70.a design (left) and a previously published WT TEM-1 crystal structure (right, PDB code 1XPB). The 70.a design contains the G251W mutation which, when introduced as a point mutation in WT TEM-1, is inactivating [1,2]. In the WT TEM-1 structure, G251 is localized to the tip of a β-strand. Two residues – F230 and P257 – occupy the vacant space adjacent to G251, effectively capping the hydrophobic core on this side of WT TEM-1. In the 70.a design, there are backbone and side chain conformational rearrangements that enable the indole side chain of W251 to cap the hydrophobic core (Supplemental Figure 11). The conformation of W251 appears to be stabilized through guanidino/indole stacking interactions with R230, which in turn forms a salt bridge with E212. In summary, alternative local interactions (W251/R230/E212) may contribute to maintaining β-lactamase structure and function.

| **Supplemental Table 1. β-Lactamase Crystallographic Data and Refinement Statistics** | | | |
|---|---|---|---|
| | 70.a | 80.a | 80.b |
| Molecule A | | | |
| Crystallization | 0.02M Divalent II, 0.1M buffer system 6, pH 8.5, 30% precipitant mix 7 (Morpheus II screen B9) | 30% PEG3350<br>0.1M MES pH 6.5 | 30% PEG3350<br>0.1M MES pH 6.5 |
| Beamline | NECAT 24ID-C | NSLS2 FMX | NSLS2 FMX |
| Wavelength (Å) | 0.97918 | 0.97932 | 0.97932 |
| Space group | P $4_1$ | P1 | P $2_1$ $2_1$ $2_1$ |
| Cell a, b, c, (Å) | 114.97, 114.97, 49.18 | 50.92, 51.12, 56.89 | 59.95, 60.9, 121.88 |
| Resolution (Å) | 45.22–3.11 (3.32-3.11) | 27.89-1.83 (1.87-1.83) | 29.57-1.59 (1.63-1.59) |
| Unique reflections | 11302 (1822) | 40574 (3405) | 58092 (5320) |
| Completeness (%) | 96.2 (87.2) | 89.95 (75.38) | 95.85 (88.77) |
| <I/> | 10.1 (3.3) | 7.30 (1.40) | 17.90 (3.40) |
| Multiplicity | 5.2 (4.6) | 2.1 (2.1) | 13.6 (12.9) |
| R-merge | 0.12 (0.418) | 0.085 (0.442) | 0.100 (0.723) |
| CC$_{1/2}$ | 0.995 (0.891) | 0.994 (0.558) | 0.999 (.887) |
| *Refinement* | | | |
| Protein residues/waters | 499/6 | 524 | 525 |
| No. reflections for R-free | 568 (57) | 1895 (162) | 1932 (184) |
| R-work | 0.218 (0.3539) | 0.1741 (0.2580) | 0.1633 (0.1831) |
| R-free | 0.2671 (0.4439) | 0.2097 (0.3195) | 0.1923 (0.2263) |
| RMSD bond lengths (Å) | 0.005 | 0.009 | 0.007 |
| RMSD bond angles (°) | 0.93 | 1.36 | 1.25 |
| Average overall *B*-factor | 68.13 | 20.73 | 16.78 |
| Mean B-factors (Å$^2$) protein/waters/Mg$^{2+}$ | 63.18/29.89 | 19.5/29.28 | 15.00/27.74 |
| Ramachandran analysis favored/allowed (%) | 90.97/8.01 | 98.46/1.54 | 98.27/1.73 |
| PDB accession code | 8OF9 | 8GII | 8GIJ |

Values in parentheses correspond to the statistics in the highest resolution bin. RMSD, root-mean-square deviation.

$R_{merge} = S_{hkl} S_j ½ I_{hkl,j} - <I_{hkl}> ½ / S_{hkl} S_{jhkl,j}$
